# N and O glycosylation of the SARS-CoV-2 spike protein

**DOI:** 10.1101/2020.07.05.187344

**Authors:** Miloslav Sanda, Lindsay Morrison, Radoslav Goldman

## Abstract

Covid-19 pandemic outbreak is the reason of the current world health crisis. The development of effective antiviral compounds and vaccines requires detailed descriptive studies of the SARS-CoV-2 proteins. The SARS-CoV-2 spike (S) protein mediates virion binding to the human cells through its interaction with the ACE2 cell surface receptor and is one of the prime immunization targets. A functional virion is composed of three S1 and three S2 subunits created by furin cleavage of the spike protein at R682, a polybasic cleavage sites that differs from the SARS-CoV spike protein of 2002. We observe that the spike protein is O-glycosylated on a threonine (T678) near the furin cleavage site occupied by core-1 and core-2 structures. In addition, we have identified eight additional O-glycopeptides on the spike glycoprotein and we confirmed that the spike protein is heavily N-glycosylated. Our recently developed LC-MS/MS methodology allowed us to identify LacdiNAc structural motives on all occupied N-glycopeptides and polyLacNAc structures on six glycopeptides of the spike protein. In conclusion, our study substantially expands the current knowledge of the spike protein’s glycosylation and enables the investigation of the influence of the O-glycosylation on its proteolytic activation.

---

The World Health Organization was informed of pneumonia cases of unknown etiology in Wuhan, Hubei Province, China on 31 December 2019 ^1^. A novel coronavirus was identified as the cause of the disease by further investigations ^2^. This new virus is related to the previously identified SARS-CoV (severe acute respiratory syndrome coronavirus) and has been named SARS-CoV-2 (severe acute respiratory syndrome coronavirus 2). Symptoms of the coronavirus disease 2019 (COVID-19) are acute onset of fever, myalgia, dyspnea, cough and evidence of ground-glass lung opacities. We do not have currently an effective vaccine or treatment for the COVID-19 patients and continued research is urgently needed to address the challenges posed by the pandemic.

Transmembrane spike (S) glycoprotein of the SARS-CoV-2 interacts with the angiotensin-converting enzyme 2 (ACE2) presented on the surface of human cells and mediates viral entry ^3–5^. Both the viral spike and the human ACE2 (hACE2) are glycoproteins and their glycosylation affects their interactions or vaccine design. Covid 19 spike glycoprotein forms a trimeric structure on the surface of the virus envelope 6. Each spike protein consists of an S1 and an S2 subunit; the S1 subunit mediates binding of the virus to the ACE2 receptor while the S2 subunit enables fusion of the virion with the cell membrane and initiates viral entry. SARS-CoV-2 has 10 to 20 times higher affinity for the ACE2 receptor than the SARS-CoV ^3^ which may be, in part, related to glycosylation of the proteins. SARS-CoV-2 S glycoprotein carries 22 N-glycosylation sequons ^6^ and at least 3 sites of mucin-type O-glycosylation were predicted ^7^ but were not yet observed experimentally. The latest analysis shows that 20 out of the 22 N-glycosylation sequons are occupied by complex, hybrid and oligomannosidic structures. Some of the sequons are predominantly occupied by oligomannose structures which could have influence on the trimeric structure. The studies also detected one O-glycopeptide occupied at sites, distinct from the predicted furin cleavage site at the S1/S2 boundary ^6,8,9^.

In this study, we report analysis of the site-specific glycoforms with focus on the resolution of structural motifs of the identified O- and N-glycopeptides. To this end, we used high-resolution LC-MS/MS with HCD fragmentation and modulated NCE ^10^ to study a recombinant SARS-CoV-2 S full-length protein expressed in human embryonic kidney (HEK 293) cells. Our analyses identified 9 occupied O-glycopeptides and 17 N-glycopeptides. We resolved, for the first time, LacdiNAc and polyLacNAc structural motifs associated with the N-glycopeptides and we identified novel O-glycopeptides including a glycopeptide near the furin cleavage site of the spike glycoprotein.

## EXPERIMENTAL SECTION

### Materials and Methods

#### Materials

Recombinant SARS-CoV-2 spike (R683A, R685A, His-tag) protein expressed in HEK 293 cell line was obtained from Acrobiosystems (Newark, DE, USA). Trypsin Gold and Glu-C, Sequencing Grade were from Promega (Madison, WI), PNGase F, Neuraminidase, 1-3 and 1-4 betagalactosidase were from New England Biolabs (Ipswich, MA).

#### Glycopeptide preparation

Aliquots of the SARS-CoV-2 S protein were dissolved in sodium bicarbonate buffer to a final concentration of 1mg/ml. The protein solution was reduced with 5 mM DTT for 60 min at 60 °C, alkylated with 15 mM iodoacetamide for 30 min in the dark, and digested with Trypsin Gold (2.5 ng/μl) at 37°C in Barocycler NEP2320 (Pressure BioSciences, South Easton, MA) for 1 hour. GluC, PNGase F, neuraminidase and beta galactosidase digests of tryptic peptides were carried out as described previously ^11,12^ with heat inactivation (99 °C for 10 min) prior to the addition of any enzyme.

#### Glycopeptide analysis using DDA nano LC-MS/MS on the Orbitrap Fusion-Lumos

Digested proteins were separated using a 120-minute ACN gradient on a 250 mm x 75 μm C18 pepmap column at a flow rate of 0.3 μL/min as described previously 13. In brief, peptide and glycopeptide separation was achieved by a 5 min trapping/washing step using 99% solvent A (2% acetonitrile, 0.1% formic acid) at 10 μL/min followed by a 90 min acetonitrile gradient at a flow rate of 0.3 μL/min: 0-3 min 2% B (0.1% formic acid in ACN), 3-5 min 2-10% B; 5-60 min 10-45% B; 60-65 min 45-98% B; 65-70 min 98% B, 70-90 min equilibration by 2% B. Glycopeptides were analyzed using Orbitrap Fusion Lumos mass spectrometer with the electrospray ionization voltage at 3 kV and the capillary temperature at 275°C. MS1 scans were performed over m/z 400–1800 with the wide quadrupole isolation on a resolution of 120,000 (m/z 200), RF Lens at 40%, intensity threshold for MS2 set to 2.0e4, selected precursors for MS2 with charge state 3-8, and dynamic exclusion 30s. Data-dependent HCD tandem mass spectra were collected with a resolution of 15,000 in the Orbitrap with fixed first mass 110 and 4 normalized collision energy 10, 20 and 35%. ETD and EThcD methods used calibrated charge dependent parameters and HCD supplemental activation was set to 15% NCE; we used the same chromatographic method and instrument settings for the ETD measurements as described above.

#### Glycopeptide analysis using cyclic ion mobility

LC-IM-MS/MS experiments were performed on a Waters Select Series cyclic ion mobility mass spectrometer with an ACQUITY M-Class solvent system. Tryptic peptides were separated using a 75 µm x 150 mm ACQUITY BEH C18 column with a 5 cm Symmetry C18 trap. Peptides were eluted over 60 minutes prior to electrospray ionization and analysis in positive mode. Glycoforms of the polybasic peptide were isolated in the quadrupole and fragmented in the trap region prior to ion mobility separations. Ion mobility methods entailing five passes of the cyclic device were previously optimized for HexNAcHex and HexNAcHexNeuAc oxonium ions and were used to separate and characterize the oxonium ion fragments of the targeted glycopeptides. Traveling wave parameters within the cyclic device were kept at default values, 375 m/s and 22V for the wave velocity and wave height, respectively. Calibration for collisional cross section was performed under a single experimental condition for both single pass and the 5-pass methods using Major Mix, with calculated uncertainties of less than 0.25% and 1%, respectively.

### Data analysis

#### Glycopeptide identification

Byonic software (protein metric) was used for the identification of summary formulas of glycans associated with the glycopeptides. Independent searches were performed on the data with different collision energy (CE) settings. All spectra of the identified glycopeptides were checked manually for the presence of structure-specific fragments. Analysis of the ion mobility data was performed using DriftScope (v2.9) by manual extraction of the retention ranges associated with the glycopeptides.

## RESULTS AND DISCUSSION

### N-glycopeptide analysis

We have identified 17 tryptic N-glycopeptides of the SARS-CoV-2 spike protein occupied by high mannose, hybrid and complex glycans. We have determined their site occupancy by PNGaseF deglycosylation in ^18^O water as described ^11^ and we find majority of the sequons fully occupied (Table 1). We have found that one sequon is not glycosylated (N603), that N234 is almost exclusively occupied by high mannose glycans. We do not have evidency for occupancy of glycosite N17. The remaining 17 sequons are dominated by complex glycans. In addition, we confirmed the presence of core fucosylation on 15 of the occupied sequons (Table 1).

**Table 1.**
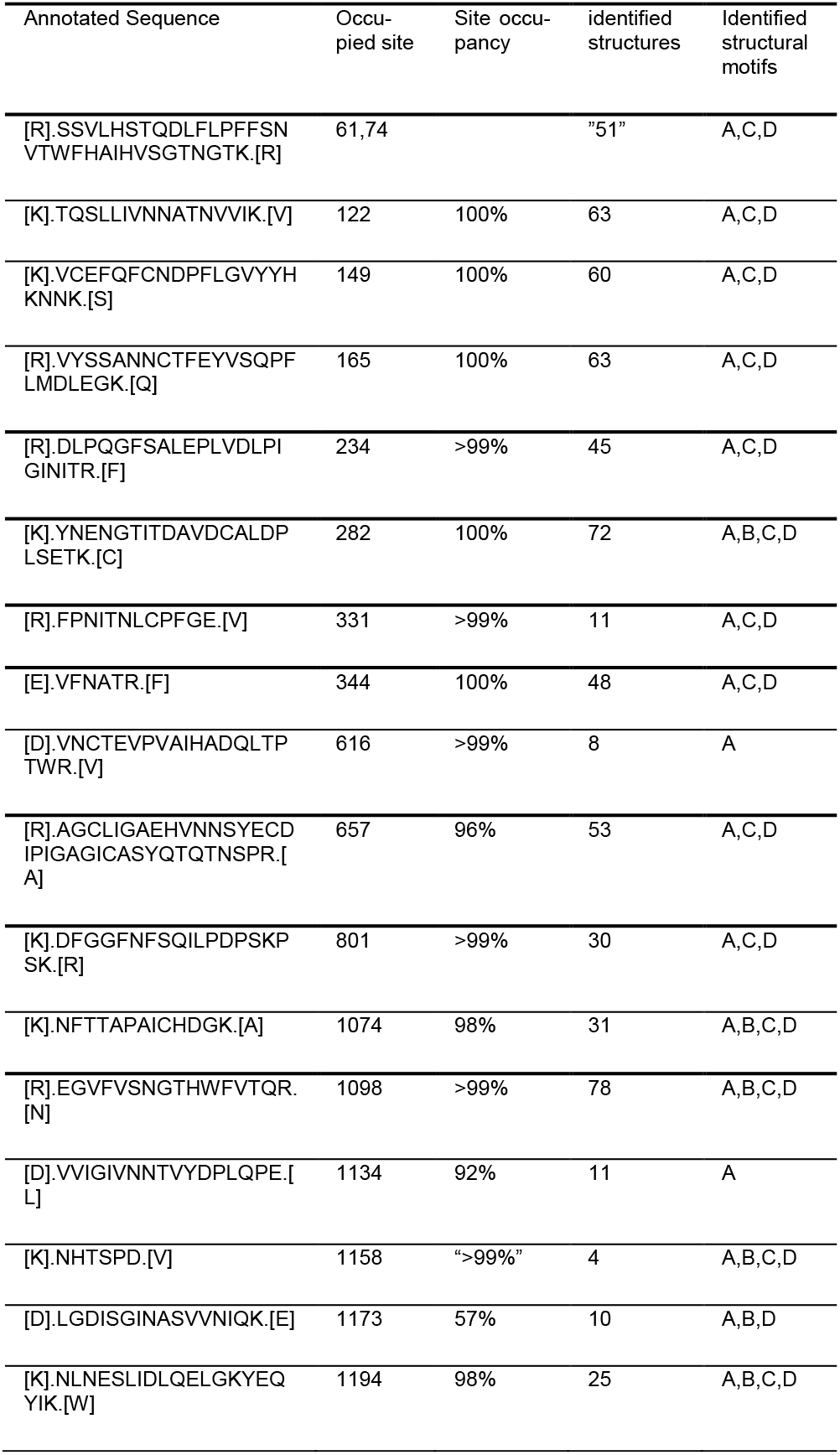
N-glycosylation of the SARS-CoV-2S glycoprotein: A LacdiNAc;B PolyLacNAc;C Outer-arm fucosylation;D core fucosylation

### Structural motifs of the N-glycans using modulated collision energy

We used our recently described workflows, using modulation of collision energy (CE) for selective fragmentation of the glycopeptides ^10^, to identify structural motifs of the N-glycosylated peptides of the SARS-CoV-2 S protein. We identified the LacdiNAc structural motif on all the occupied sequons of the SARS-CoV-2 S expressed in the HEK293 cells (Table 1). The low CE tandem mass spectrum (Figure 1) reveals structural features of an asymmetric LacdiNAc motif contained within a disialylated biantennary N-glycan. Presence of the m/z 366/407 ions distinguishes the LacNAc and LacdiNAc motifs; in addition, we observe the m/z 657/698 ions of their sialylated counterparts. In addition to the fucosylated and/or sialylated LacdiNAc, we also identified polyLacNAc structures on 5 N-glycopeptides (Table 1) and we resolved extensive fucosylation of the core as well as the outer-arms of the N-glycopeptides as described previously ^10^. The presence of core fucose was confirmed on 15 sequons and we confirmed the presence of outer arm fucosylation ^11,14^ on 7 sequons which is in contrast to the previously published data ^8^. This might be a result of slight differences in the HEK293 expression systems used or differences in the analytical methods. For example, our study analyzed a modified full-length protein not cleaved by convertases which could potentially cause some differences. It is, however, more likely that the energy optimized workflows improve the structural resolution. We do not achieve complete assignment of all linkages or quantification of the isobaric structures but the presence of these structural motifs, frequently associated with specific biological functions, is clearly established. The overall results show that 5 glycopeptides carry polyLacNac motifs, that all sequons occupied by complex glycans carry LacdiNAc to some degree, and that the LacdiNAc structures constitute majority (>50%) of the glycoforms on N165 and N1098. This may not necessarily reflect the N-glycoforms of a virion but the HEK293 expression system is commonly used for functional studies of the S glycoprotein or the production of vaccine candidates which means that resolution of the structures is highly relevant.

**Figure 1.**
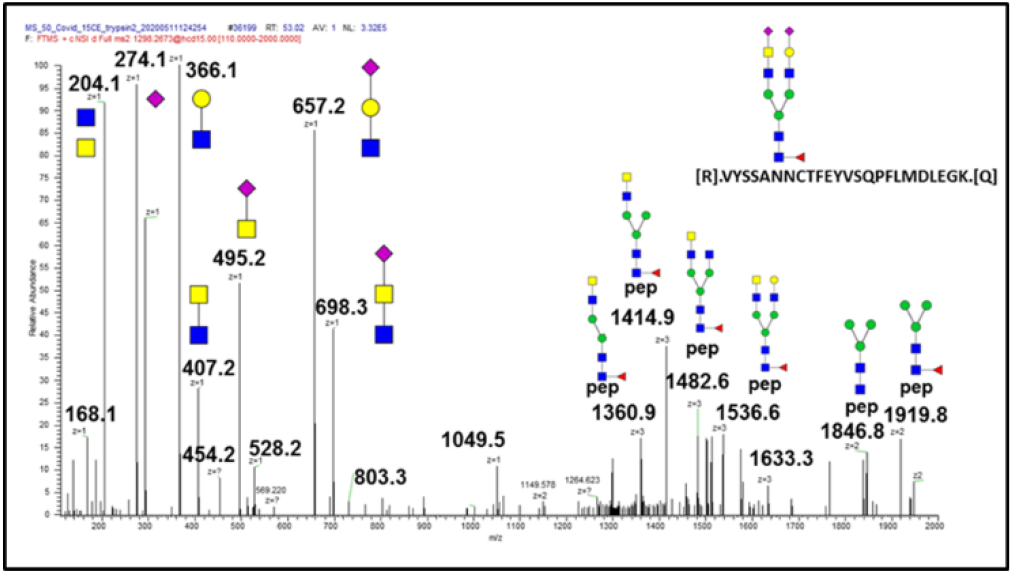
HCD fragmentation of the N165 glycopeptide carrying an asymmetric biantennary glycan with sialylated LacdiNAc structural motif.

### O-glycopeptide analysis

Previously published data describes one O-glycopeptide occupied at S323 and T325 6,8. We identified the same O-glycopeptides but, in addition, we have identified 8 O-glycopeptides occupied by core-1 and core-2 structures (Table 2 and supplemental table 1). Occupancy of the sites varies from <1% to 57% and is very low for at least three of the glycopeptides. However, we detect approximately 13% occupancy with core-1 and core-2 structures at the T678 (Figure 2 and 3) located near the polybasic furin cleavage site between the S1 and S2 subunits which evolved in the SARS-CoV-2 S protein ^7^. This is relevant because O-glycans proximal to the convertase cleavage sites of protein substrates regulate proteolysis ^15^ and may regulate activation of the SARS-CoV-2 S protein. Supplemental Figure 1 documents identification of an O-glycopeptide following deglycosylation with PNGaseF. This is interesting because O-glycosylation in such a close proximity to N-glycosylation is rarely described; we do not know if this has any functional relevance but it shows that analysis of N-deglycosylated peptides for O-glycoforms may deserve attention. Retention times of the T687 O-glycoforms (Figure 4) follow the expected trends of structure dependent reverse phase chromatographic behavior of glycopeptides ^16,17^.

**Table 2.**
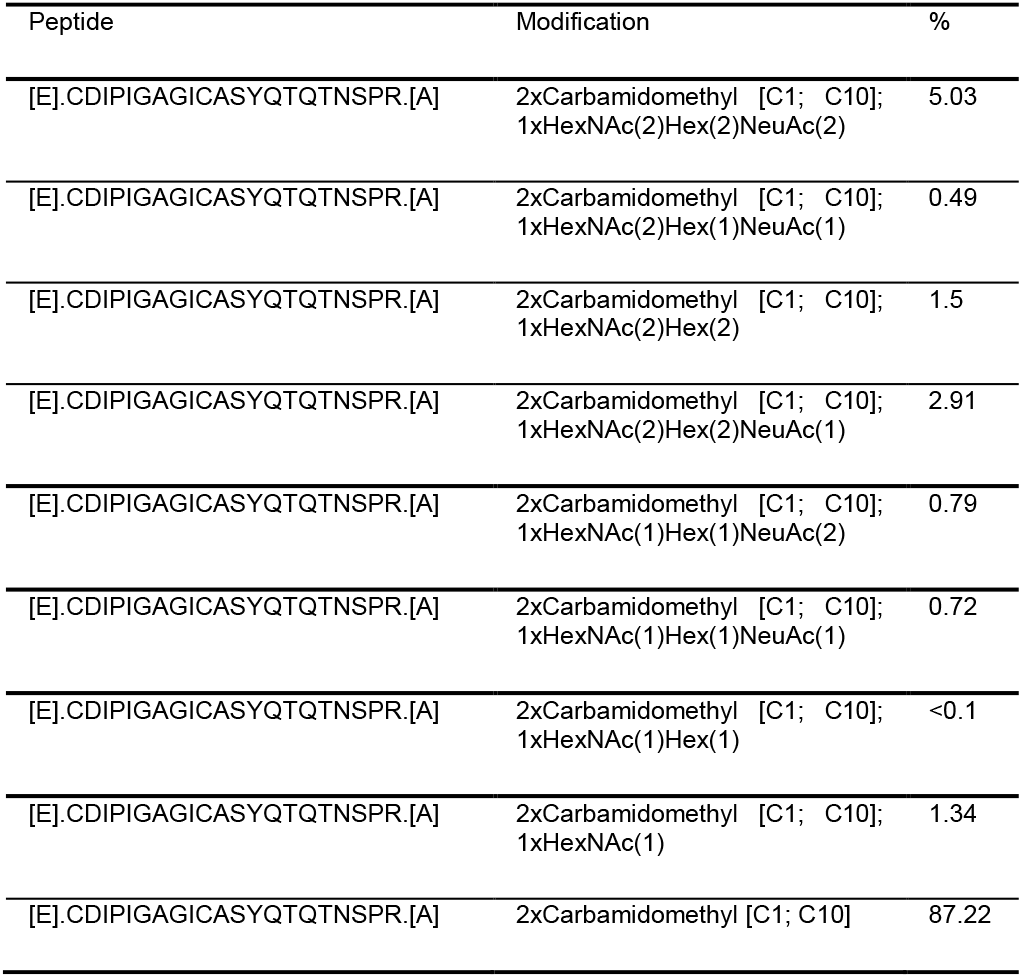
O-glycosylation of the SARS-CoV-2 S glycoprotein on glycosite T678

**Figure 2.**
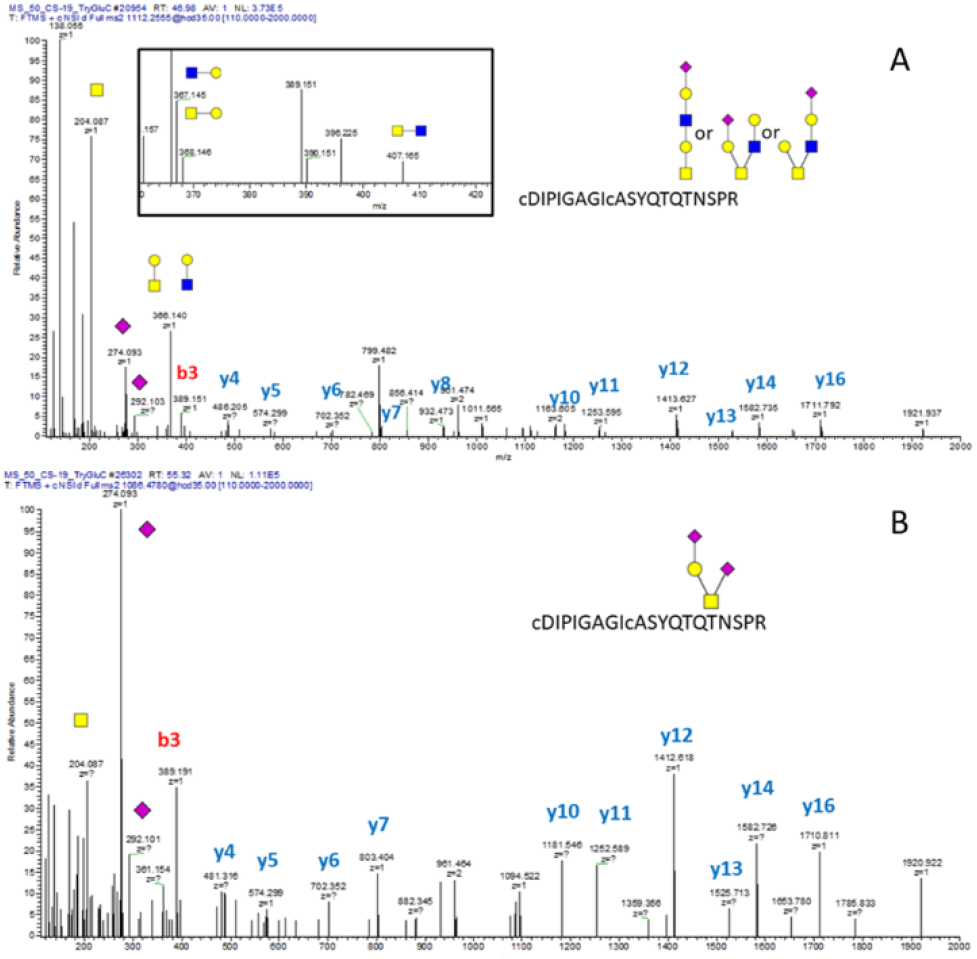
HCD tandem mass spectra of the SARS-CoV-2 S protein O-glycosylated on T678 with the following structures: (A) extended core-1 and core-2 structures; (B) disialylated core-1 structure. Inset: oxonium ions in the HCD fragmentation spectrum confirm the presence of core-2 structures.

**Figure 4.**
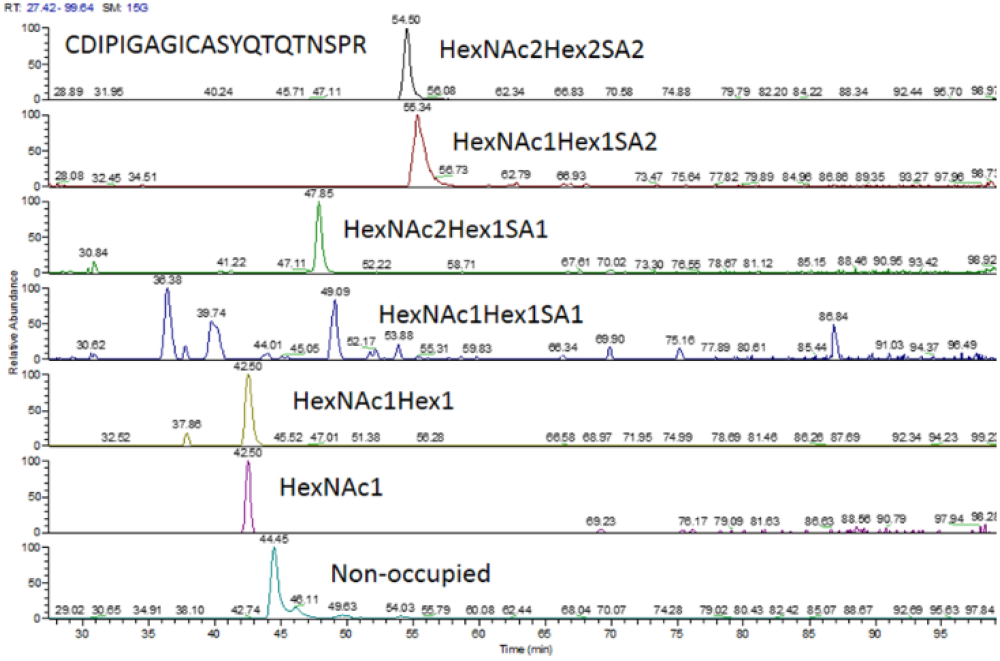
The CDIPIGAGICASYQTQTNSPR O-glycopeptides of SARS-CoV-2 S protein with the expected glycoform-dependent RT shifts visible in the XIC chromatograms.

**Figure 5.**
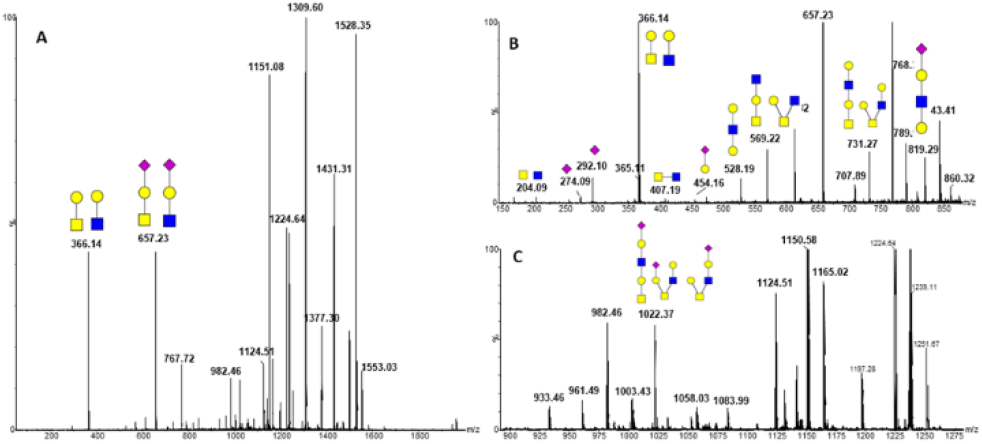
Beam type tandem mass spectra of the AGC(cam)LIGAEHVNN(dea)SYEC(cam)DIPIGAGIC(cam)ASYQTQTNSPR (HexNAc2Hex2SA1) O-glycopeptide with assigned extended core-1 and core-2 structures. The structures are characterized by the following fragments: (A) oxonium ions 366 and 657, generated from both core-1 and core-2 structures; (B) oxonium ion 407 specific for the core-2 and ions 528 and 819 specific to the extended core-1 structure; and (C) oxonium ion 1022 correspoding to the detached intact glycans.

glycoprotein. We choose to use cIMS on the fragment to reduce influence of the peptide backbone on the structural resolution. We were able to confirm the presence of core-2 structures by the diHexNAc fragment m/z 407 in the HCD spectra using the Orbitrap Fusion Lumos (Figure 2A inset). The tandem mass spectra obtained from the cIMS instrument preserve large oxonium ions, such as the intact detached glycan m/z 1022 (Figure 5) which confirms that a hexasacharide occupies the O-glycopeptide AGC(cam)LIGAEHVNN(dea)SYEC(cam)DIPIGAGIC(cam)ASYQTQTNSPR but using beam type fragmentation we could not determine which serine or threonine is occupied. We cannot fully exclude the possibility of contribution from a second glycan at this peptide but neither the ETD not the HCD spectra show evidence of another occupied site besides the T678 of this peptide. We have also confirmed the presence of an extended core-1 structure associated with this glycopeptide by the fragments 528 and 819 observed in the spectra (Figure 5B).

### Structural analysis of the O-glycopeptides using cIMS

We have used cIMS to separate isomeric oxonium ion fragments of the glycopeptides. We have used the m/z 657 ion to assign sialylation of the core-2 monosialylated structures. We used an optimized procedure based on a hemopexin glycopeptide standard, which we described previously ^12^, and we determined CCS of the fragment 657 (Figure 6) observed by fragmentation of the glycopeptide with sialyl-T antigen with linkage (α2-3) (CCS 234.9) and by fragmentation of an N-glycopeptide with sialyl-LacNAc with (α2-6) linkage (CCS 232.8) (data not shown).

**Figure 6.**
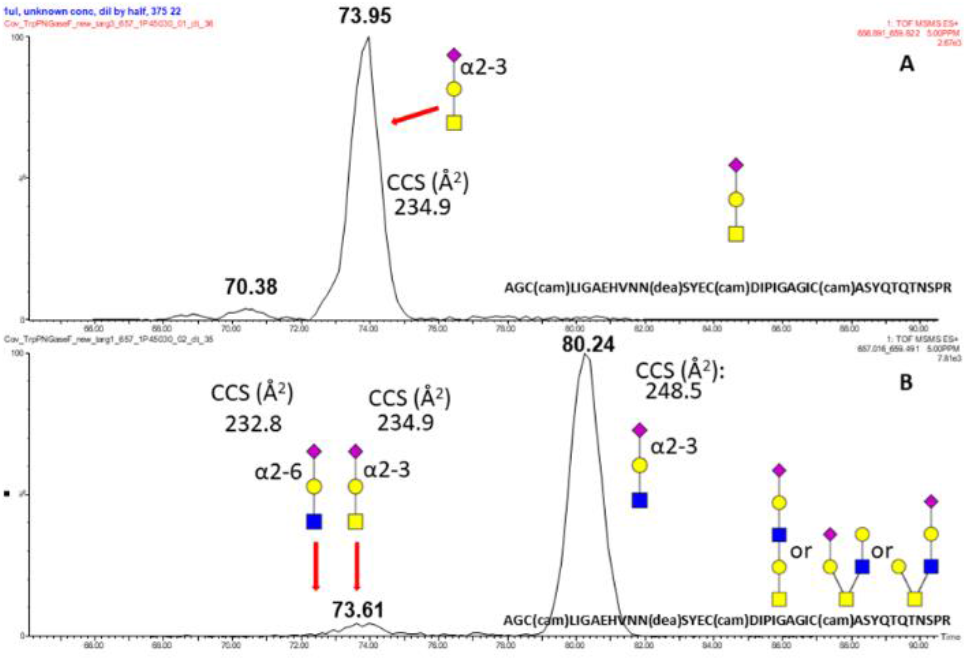
cIMS of the fragment m/z 657 with measured CCS assignments produced by fragmentation of the AGC(cam)LIGAEHVNN(dea)SYEC(cam)DIPIGAGIC(cam)ASYQTQTNSPR (HexNAcHexSA) (A) and (HexNAc2Hex2SA) (B) O-glycopeptides produced by tryptic digests and PNGaseF deglycosylation of the SARS-CoV-2 S glycoprotein

This is in agreement with the previously published results on the linkages of the sialylated glycans ^6,8^. We resolved two major IMS peaks in the cIMS of the fragment m/z 657 using a one pass method (Figure 7A). With 5 passes, the first peak was partially separated into two analytes with determined CCSs of 232.8 and 234.9 and a second peak CCS 248.5. This is reproducible for all 2HexNAc containing structures (Supplemental Figure 2). The CCS of the first peak fits exactly the previously observed CCS of sialylated α2-6 LacNAc while the CCS of the second peak fits the CCS of the silylated α2-3 T-antigen. CCS 248.5 of the third peak is in agreement with the previously described CCS of α2-3 LacNAc ^19 20^. The peak corresponding to the 2-6 linked sialic acid is better visible under high collision energy (Figure 7B-D) due to different stability of the SA-Gal bond ^21^. We have determined a 7/3 ratio of the GlcNAc-Gal-2-3-SA and GalNAc-Gal-2-3SA in the mono-sialylated core-2 structure (Figure 7, panel A). We used the cIMS of the fragment m/z 731 (2HexNAc2Hex) to determine the ratio of the core-2 structure and the extended core-1 structure. We obtained 2 major peaks using 5 passes of the cIMS (data not shown); the first peak (time:69.70; CCS: 238.8) is consistent with a core-2 structure and the second peak (76.84; CCS: 251.5) with a linear core-1 extended structure with terminal GalNAc(1-3)Gal as described previously ^22^. Ratio of the core-1 with terminal GalNAc(1-3)Gal and the the core-2 structure is 25/75.

**Figure 7.**
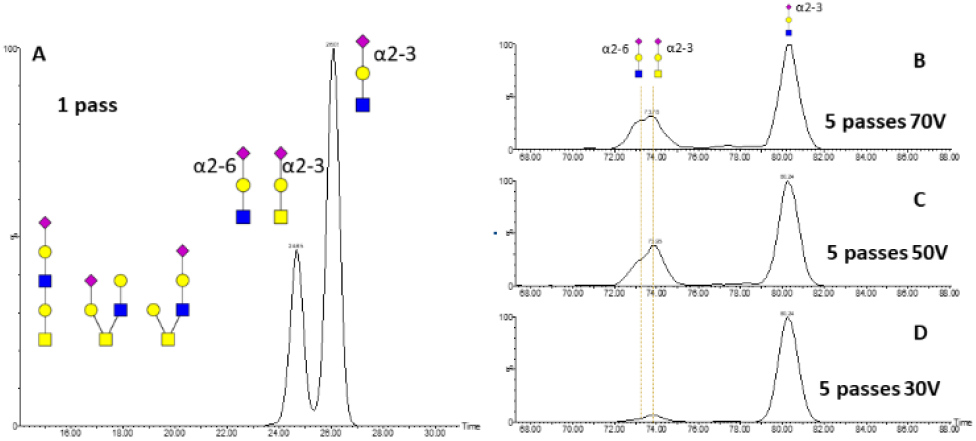
cIMS of the m/z 657 fragment produced by fragmentation of the AGC(cam)LIGAEHVNN(dea)SYEC(cam)DIPIGAGIC(cam)ASYQTQTNSPR O-Glycopeptide produced by tryptic digest and PNGaseF deglycosylation of the SARS-CoV-2 S glycoprotein using the following settings: (A) one pass cIMS does not resolve GalNAcGal(2-3)SA and GlcNAcGal(2-6)SA; (B). 5 passes cIMS with 70V CE; (C) 5 passes cIMS with 70V CE; and (D) 5 passes cIMS with 70V CE. The ion mobilograms (B,C,D) show that multiple passes improve resolution of isobaric (GalNAcGal(2-3)SA and GlcNAcGal(2-6)SA) structures and reveals differences in the stability of the sialic acid linkages.

## CONCLUSION

We have used our energy optimized LC-MS/MS and ion mobility MS/MS methods to resolve structural motifs of the N- and O-glycans of the SARS-CoV-2 S protein. We identified 17 N-glycopeptides, with many glycoforms including the LAcdiNAc and polyLacNAc structural motives. This is important for functional studies and the use of the protein as an immunization target. In addition, we identified, for the first time, an O-glycopeptide adjacent to the polybasic furin cleavage site located between the S1/S2 subunits that carries core-1 and core-2 structures capped primarily with α2-3 sialic acid at the T678. The furin cleavage site is unique to the SARS-CoV-2 S protein compared to the SARS-CoV of 2002 and its cleavage is potentially regulated by the nearby O-glycans as described for other convertases. In addition, we identified 8 additional O-glycopeptides of variable occupancy and unknown functional significance. The study expands substantially the knowledge of the glycoforms of SARS-CoV-2 S expressed in the HEK293 cells and warrants further exploration of the impact of glycosylation on the S protein’s function

## ASSOCIATED CONTENT

### Author Contributions

All the authors contributed to the writing of the manuscript and gave approval to the final version of the manuscript.

### Notes

The authors declare no competing financial interest.

## ACKNOWLEDGMENT

Research reported in this publication was supported by the National Institutes of Health under awards S10OD023557, U01CA230692, and R01CA238455 to RG. The content is solely the responsibility of the authors and does not necessarily represent the official views of the National Institutes of Health.

