## Supplemental Material for "N and O glycosylation of the SARS-CoV-2 spike protein"

Supplemental informations

1. Supplemental Table 1.: O-glycosylation of the SARS-CoV-2 S glycoprotein

2. Supplemental Figure 1: HCD tandem mass spectrum of the O-glycopeptide YFKNHTSPDVD deglycosylated with PNGaseF shows that the threonine or serine are occupied by a core-2 glycan. This peptide was occupied by N- and O-glycans in neighboring positions prior to the deglycosylation with PNGaseF.

3. Supplemental Figure 2: cIMS ionmobilograms of the 657 fragment derived from the fragmentation of the AGC(cam)LIGAEHVNN(dea)SYEC(cam)DIPIGAGIC(cam)ASYQTQTNSPR O-Glycopeptide treated with PNGaseF and occupied by different O-glycan structures as indicated.

4. Supplemental Figure 3: HCD spectra of the trypsin/GluC O-glycopeptide CDIPIGAGICASYQTQT(HexNAc)NSPR occupied by Tn-antigen following treatment with neuraminidase, beta galactosidase, and PNGaseF.

Supplemental Table 1. O-glycosylation of the SARS-CoV-2 S glycoprotein

| Peptide | Modification | % |
|---|---|---|
| [E].CDIPIGAGICASYQTQTNSPR.[A] | 2xCarbamidomethyl [C1; C10]; 1xHexNAc(2)Hex(2)NeuAc(2) | 5.03 |
| [E].CDIPIGAGICASYQTQTNSPR.[A] | 2xCarbamidomethyl [C1; C10]; 1xHexNAc(2)Hex(1)NeuAc(1) | 0.49 |
| [E].CDIPIGAGICASYQTQTNSPR.[A] | 2xCarbamidomethyl [C1; C10]; 1xHexNAc(2)Hex(2) | 1.5 |
| [E].CDIPIGAGICASYQTQTNSPR.[A] | 2xCarbamidomethyl [C1; C10]; 1xHexNAc(2)Hex(2)NeuAc(1) | 2.91 |
| [E].CDIPIGAGICASYQTQTNSPR.[A] | 2xCarbamidomethyl [C1; C10]; 1xHexNAc(1)Hex(1)NeuAc(2) | 0.79 |
| [E].CDIPIGAGICASYQTQTNSPR.[A] | 2xCarbamidomethyl [C1; C10]; 1xHexNAc(1)Hex(1)NeuAc(1) | 0.72 |
| [E].CDIPIGAGICASYQTQTNSPR.[A] | 2xCarbamidomethyl [C1; C10]; 1xHexNAc(1)Hex(1) | <0.1 |
| [E].CDIPIGAGICASYQTQTNSPR.[A] | 2xCarbamidomethyl [C1; C10]; 1xHexNAc(1) | 1.34 |
| [E].CDIPIGAGICASYQTQTNSPR.[A] | 2xCarbamidomethyl [C1; C10] | 87.22 |
| [R].TQLPPAYTNSFTR.[G] | | 99.73 |
| [R].TQLPPAYTNSFTR.[G] | 1xDeamidated [N9] | 0.18 |
| [R].TQLPPAYTNSFTR.[G] | 1xHexNAc(1) | 0.05 |
| [R].TQLPPAYTNSFTR.[G] | 1xHexNAc(1)Hex(1) | 0.02 |
| [R].TQLPPAYTNSFTR.[G] | 1xHexNAc(3) | 0.01 |
| [R].TQLPPAYTNSFTR.[G] | 1xHexNAc(1)Hex(1)NeuAc(2) | 0.01 |
| [K].YFKNHTSPDVD.[L] | 1xDeamidated [N4]; 1xHexNAc(1)Hex(1)NeuAc(2) | 7.18 |
| [K].YFKNHTSPDVD.[L] | 1xDeamidated [N4] | 70.33 |
| [K].YFKNHTSPDVD.[L] | 1xDeamidated [N4]; 1xHexNAc(2)Hex(2)NeuAc(2) | 22.49 |
| [R].VQPTESIVR.[F] | 1xHexNAc(2)Hex(2)NeuAc(1) | 7.13 |
| [R].VQPTESIVR.[F] | 1xHexNAc(1)Hex(1)NeuAc(2) | 30.56 |
| [R].VQPTESIVR.[F] | 1xHexNAc(1)Hex(1)NeuAc(1) | 5.65 |
| [R].VQPTESIVR.[F] | 1xHexNAc(1)Hex(1) | 13.93 |
| [R].VQPTESIVR.[F] | | 42.75 |
| [R].VYSTGSNVFQTR.[A] | 1xHexNAc(1) | 0.023 |
| [R].VYSTGSNVFQTR.[A] | | 99.67 |
| [E].VPVAIHADQLTPTWR.[V] | 1xHexNAc(1)Hex(1)NeuAc(2) | 0.013 |
| [E].VPVAIHADQLTPTWR.[V] | | 99.98 |
| [D].STECSNLLLQYGSFCTQLNR.[A] | 2xCarbamidomethyl [C4; C15]; 1xHexNAc(1)Hex(1); 1xHexNAc(2)Hex(3)Fuc(1) | 87.23 |
| [D].STECSNLLLQYGSFCTQLNR.[A] | 2xCarbamidomethyl [C4; C15]; 1xHexNAc(2) [S/T]; 1xHexNAc(2)Hex(1)NeuAc(2) | 12.77 |
| [K].LPDDFTGCVIAWNSNNLD.[S] | 1xCarbamidomethyl [C8] | 82.04 |
| [K].LPDDFTGCVIAWNSNNLD.[S] | 1xCarbamidomethyl [C8]; 2xHexNAc(1)Hex(2)Fuc(1) | 14.37 |
| [K].LPDDFTGCVIAWNSNNLD.[S] | 1xCarbamidomethyl [C8]; 1xHexNAc(3)Hex(2)Fuc(1)NeuAc(1) | 3.59 |
| [D].YSVLYNSASFSTFK.[C] | 1xDeamidated [N6] | 3.35 |
| [D].YSVLYNSASFSTFK.[C] | 1xDeamidated [N6]; 1xHexNAc(1)Hex(1)Fuc(1)NeuAc(1) | 0.037 |
| [D].YSVLYNSASFSTFK.[C] | | 96.61 |

Supplemental Figure 1. HCD tandem mass spectrum of the O-glycopeptide YFKNHTSPDVD deglycosylated with PNGaseF shows that the threonine or serine are occupied by a core-2 glycan. This peptide was occupied by N- and O-glycans in neighboring positions prior to the deglycosylation with PNGaseF.

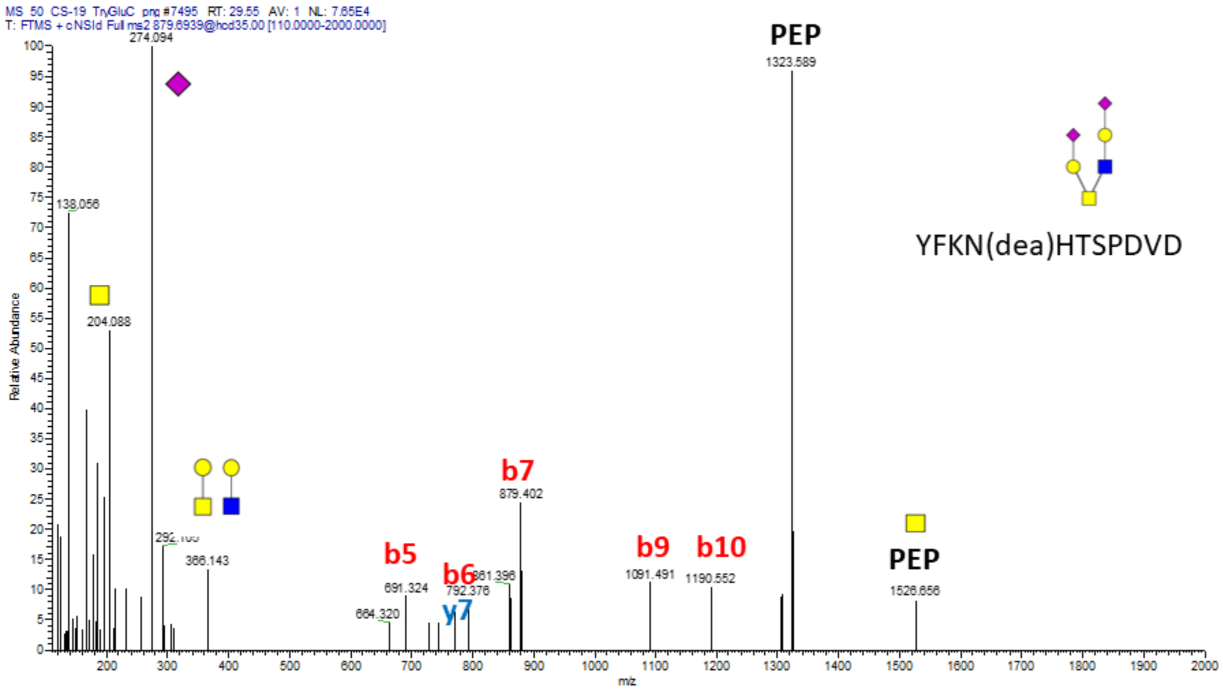

Supplemental Figure 2. cIMS ionmobilograms of the 657 fragment derived from the fragmentation of the AGC(cam)LIGAEHVNN(dea)SYEC(cam)DIPIGAGIC(cam)ASYQTQTNSPR O-Glycopeptide treated with PNGaseF and occupied by different O-glycan structures as indicated.

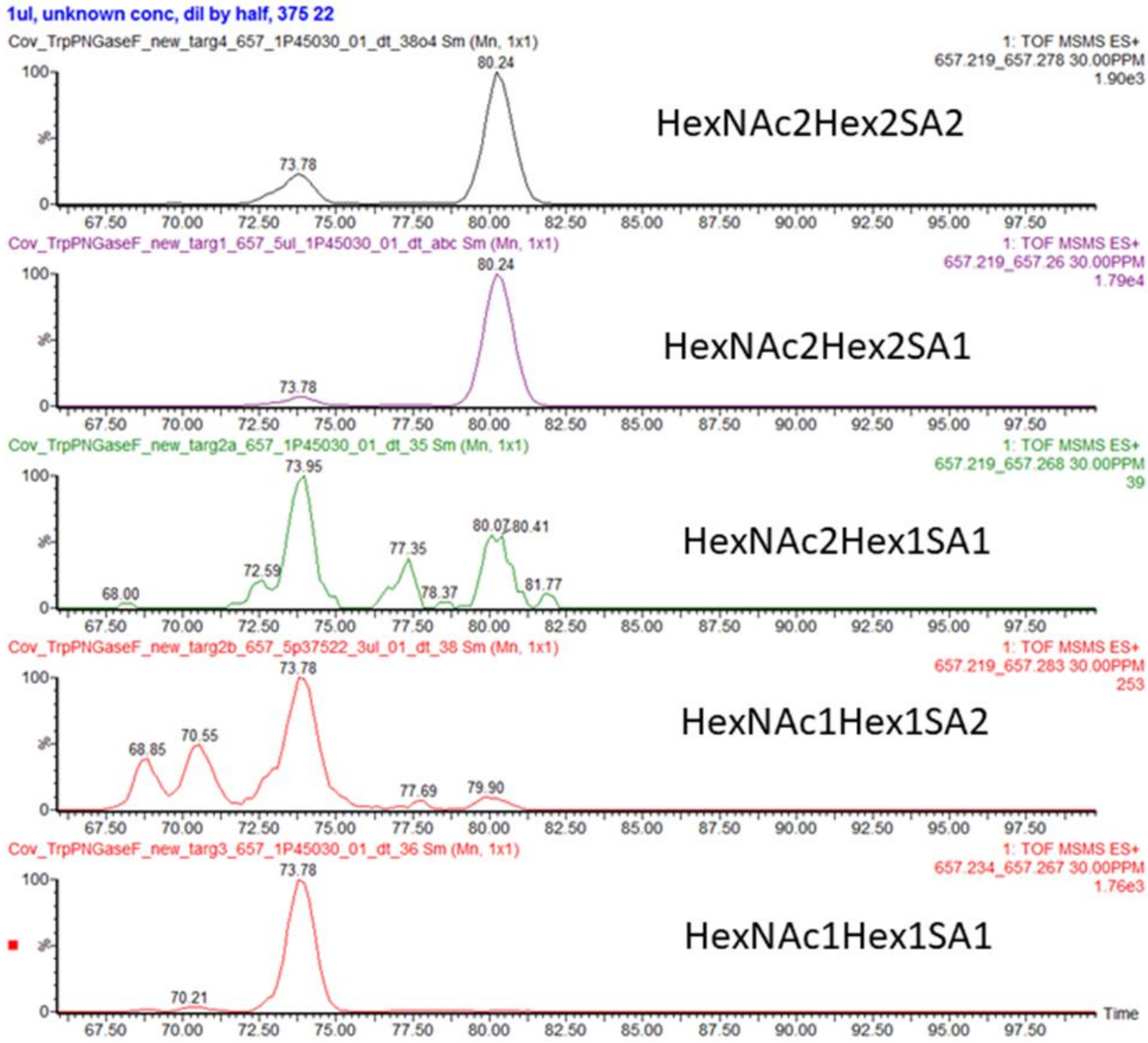

Supplemental Figure 3. HCD spectra of the trypsin/GluC O-glycopeptide CDIPIGAGICASYQTQT(HexNAc)NSPR occupied by Tn-antigen following treatment with neuraminidase, beta galactosidase, and PNGaseF.

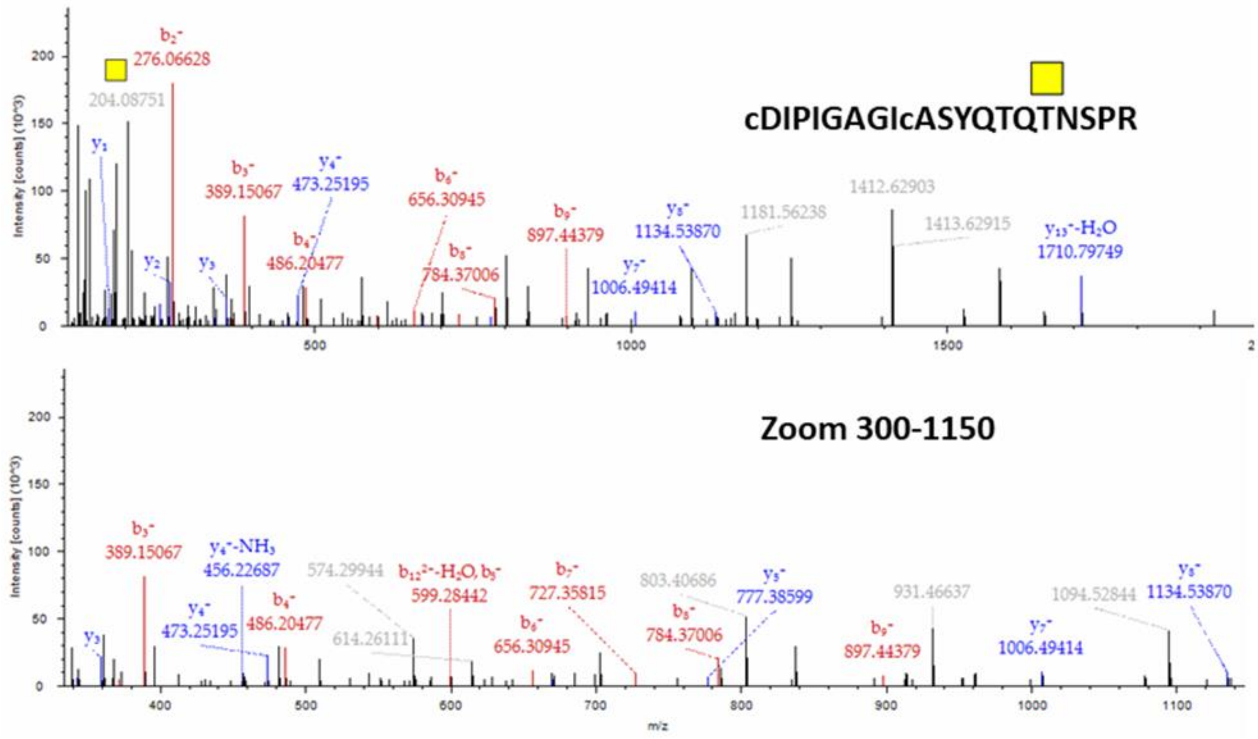
